# A spiking neural network model for proprioception of limb kinematics in insect locomotion

**DOI:** 10.1101/2024.09.27.615365

**Authors:** Thomas van der Veen, Yonathan Cohen, Elisabetta Chicca, Volker Dürr

## Abstract

Proprioception plays a key role in all behaviours that involve the control of force, posture or movement. Computationally, many proprioceptive afferents share three common features: First, their strictly local encoding of stimulus magnitudes leads to range fractionation in sensory arrays. As a result, encoding of large joint angle ranges requires integration of convergent afferent information by first-order interneurons. Second, their phasic-tonic response properties lead to fractional encoding of the fundamental sensory magnitude and its derivatives (e.g., joint angle and angular velocity). Third, the distribution of disjunct sensory arrays across the body accounts for distributed encoding of complex movements, e.g., at multiple joints or by multiple limbs. The present study models the distributed encoding of limb kinematics, proposing a multi-layer spiking neural network for distributed computation of whole-body posture and movement. Spiking neuron models are biologically plausible because they link the sub-threshold state of neurons to the timing of spike events. The encoding properties of each network layer are evaluated with experimental data on whole-body kinematics of unrestrained walking and climbing stick insects, comprising concurrent joint angle time courses of 6 × 3 leg joints. The first part of the study models strictly local, phasic-tonic encoding of joint angle by proprioceptive hair field afferents by use of Adaptive Exponential Integrate-and-Fire neurons. Convergent afferent information is then integrated by two types of first-order interneurons, modelled as Leaky Integrate-and-Fire neurons, tuned to encode either joint position or velocity across the entire working range with high accuracy. As in known velocity-encoding antennal mechanosensory interneurons, spike rate increases linearly with angular velocity. Building on distributed position/velocity encoding, the second part of the study introduces second- and third-order interneurons. We demonstrate that simple combinations of two or three position/velocity inputs from disjunct arrays can encode high-order movement information about step cycle phases and converge to encode overall body posture.

**Author summary:** When stick insects climb through a bramble bush at night, they successfully navigate through highly complex terrain with little more sensory information than touch and proprioception of their own body posture and movement. To achieve this, their central nervous system needs to monitor the position and motion of all limbs, and infer information about whole-body movement from integration in a multi-layer neural network. Although the encoding properties of some proprioceptive inputs to this network are known, the integration and processing of distributed proprioceptive information is poorly understood. Here, we use a computational model of a spiking neural network to simulate peripheral encoding of 6 × 3 joint angles and angular velocities. The second part of the study explores how higher-order information can be integrated across multiple joints and limbs. For evaluation, we use experimental data from unrestrained walking and climbing stick insects. Spiking neurons model the key response properties known from their real biological counterparts. In particular, we show that the first integration layer of the model is able to encode joint angle and velocity both linearly and accurately from an array of phasic-tonic input elements. The model is simple, accurate and based, where possible, on biological evidence.

## Introduction

Proprioception is the ability to perceive body posture, movement, and load, a ubiquitous ‘sixth sense’ in all mobile organisms [1]. It relies on information provided by mechanosensory neurons, known as proprioceptors. In insects and other arthropods, proprioceptors are distributed throughout the entire musculo-skeletal system, most of them embedded in or attached to the cuticle [2]. Multiple types of mechanoreceptors encode physical magnitudes as different as force, joint angle, or angular velocity [3, 4], with a high degree of convergence of afferent information from distinct sensors [5–7]. Various proprioceptor types are distributed across the body (e.g., hair fields: [8]; chordotonal organs: [9]) or leg parts (e.g., campaniform sensilla: [10, 11]), with distinct arrangements of sensory arrays on single limb segments (e.g., in campaniform sensilla: [12]) or at joints (e.g., in hair fields: [13]). In the context of insect locomotion, proprioception plays a crucial role in regulating reflexes, ensuring precise phase timing and controlling the activation timing of specific muscles [1, 14, 15], but also in spatial coordination of the legs and antennae [16]. A key aspect for these functions is the representation of the position and movement of the limb within central nervous system (CNS). While several types of interneurons (INs) that encode limb movements based on proprioceptive signals have been identified for legs [17, 18] and antennae [19, 20], and connectomics research promises the mapping of entire processing networks [21], the computational aspects of proprioceptive information processing are still poorly understood.

Here, we propose a multi-layer spiking neural network (SNN) for processing of distributed proprioceptive information about limb kinematics. The main objective is to encode increasingly complex information about posture and movement in a feed-forward network. We thus neglect the use of proprioceptive information in feed-back control, focusing on internal representation and state estimation instead [22]. To this end, the first part of our study models the phasic-tonic spike response of hair field afferents, and convergent encoding of position and velocity in first-order INs. The second part then focuses on the integration of distributed proprioceptive information from multiple joints per leg, and from multiple legs to encode whole-body movement (van der Veen et al., companion paper [23]). We use SNNs [24] for their similarity to biological neurons, integrating neural and synaptic states, temporal dynamics, and generation of action potentials.

This first part of the study addresses two major computational challenges of peripheral proprioceptive encoding. The first challenge concerns the phasic-tonic spike frequency response to step changes in the stimulus magnitude. All of the main proprioceptors in insects, i.e., campaniform sensilla [25], chordotonal organs [9], and hair fields [26], display a mix of tonic (slowly adapting) and phasic (rapidly adapting) response components to step changes in load [27], joint angle or angular velocity [1, 28]. This is caused both by the mechanics of the surrounding or embedding structure [29], and by the properties of the sensory neuron [30]. As a consequence, the afferent spike rate encodes the input magnitude itself (e.g., joint angle) along with its derivatives (e.g., angular velocity). Models of specific sensorimotor pathways have exploited this dynamic encoding for computational purposes in individual INs [31] or for PD control of steering [32], yet others have focused on cellular mechanisms underlying spike rate adaptation [30] or have devised compact mathematical descriptions thereof [33]. In a similar non-spiking model, Ache and Dürr (2015) [34] utilized a lead-lag-system of parallel filter blocks to simulate phasic-tonic spike rate dynamics. Our present model takes an intermediate approach, employing the mathematically compact Adaptive Exponential Integrate-and-Fire (AdEx) [35] for rate adaptation of a spiking neuron.

The second computational challenge concerns the fact that, quite generally, mechanotransduction encodes local forces and/or deformations. Therefore, a common feature of all proprioceptive afferents is their strictly local encoding of sensory magnitudes. As a consequence, proprioceptive organs are sensory arrays that show range fractionation of their overall receptive field [36]. Computationally, this requires convergence of multiple afferents to obtain a first-order IN with a suitably large receptive field. The present study addresses the problem of how a first-order IN may be tuned to either position or velocity only, despite mixed encoding of position and velocity in its input elements [19]. As a particular example, we attempt to model linear velocity encoding across a wide range of input velocities, as described for first-order INs of the antennal mechanosensory pathway [37]. Other than an earlier model that used continuous filter functions for parallel encoding of joint angle and angular velocity [34], our present model uses a simple Leaky Integrate-and-Fire (LIF) neuron model along with appropriate arrangement of the sensory input array.

Our SNN was tuned and evaluated in conjunction with electrophysiological data on proprioceptive hair fields (the scapal hair plate of the American cockroach *Periplaneta americana* [38]) and motion capture data on whole-body kinematics of unrestrained walking and climbing stick insects *Carausius morosus* [39]. Stick insects bear proprioceptive hair fields at all limb bases (antennae: [13]; legs: [26, 40]; Fig 1A) where they are located on the cuticle near joint membranes (Fig 1B). When deflected, the hair acts as a lever arm, exerting a force on a sensory neuron dendrite [3], typically causing a phasic-tonic response [28, 38]. In stick insects, ablation of individual hair fields affects leg positioning [41, 42], altered swing height [43] or antennal inter-joint coordination [13], underscoring their relevance in the control of natural locomotion behaviour.

**Fig 1.**
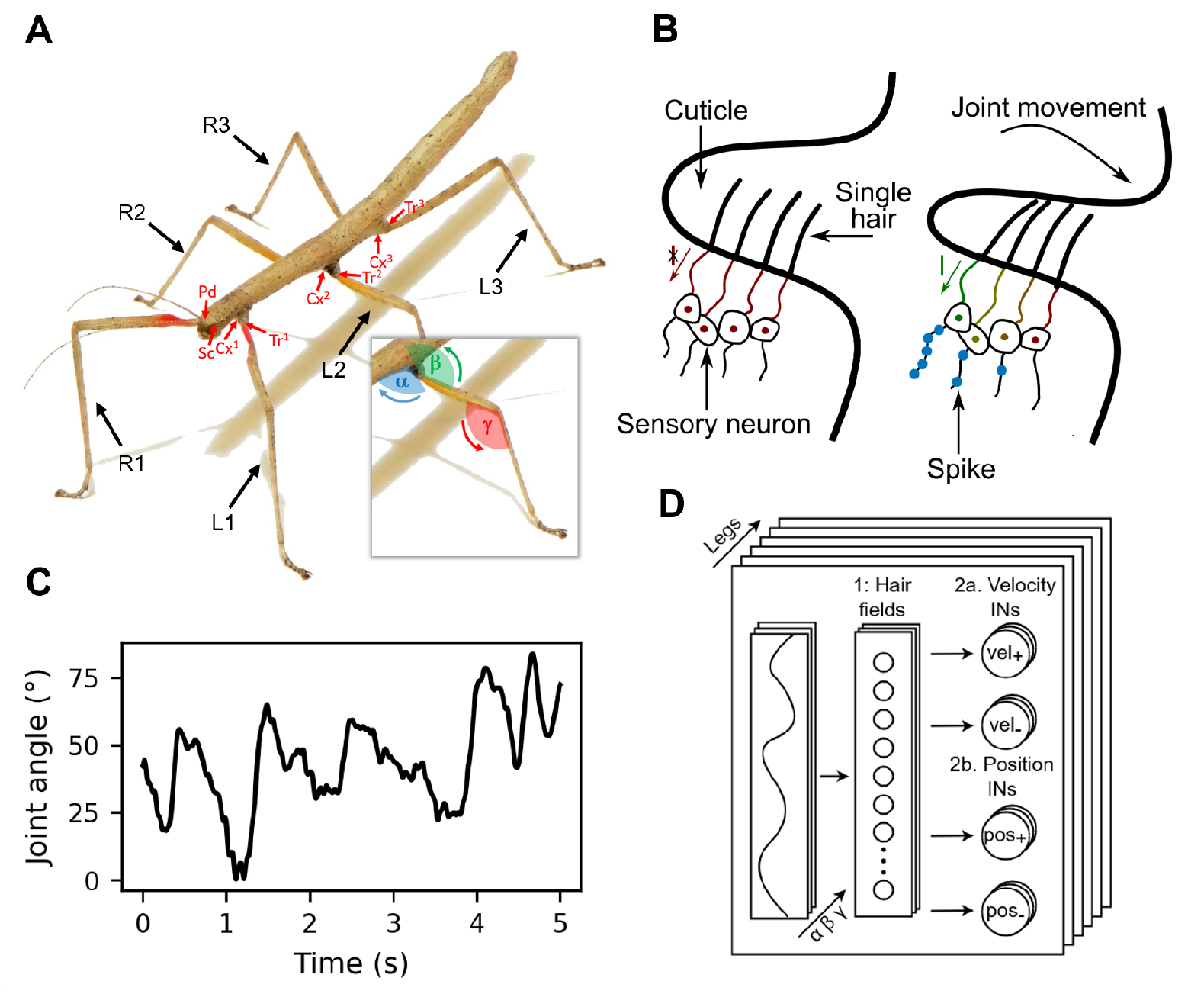
Distributed proprioceptive encoding of leg posture and movement. **A**. The six legs of the Indian stick insect *Carausius morosus* have similar size and structure. The dataset used here includes joint angles of the thorax-coxa, coxa-trochanter and femur-tibia joints, referred to as the *α, β* and *γ* joints, respectively. Red arrows indicate locations of proprioceptive hair fields at the *α* and *β* joints of the legs (labeled Cx and Tr, respectively) as well as the head-scape and scape-pedicel joints on the antenna (labeled Sc and Pd, respectively). **B**. Schematic representation of a proprioceptive hair field on the cuticle near a joint, before and during joint movement. A change in joint angle causes hair deflection. As the joint angle increases, the number of hairs deflected and the deflection angle per hair also increase. Each hair acts as a lever arm connected to a dendrite of a sensory neuron. Mechanotransduction channels open, allowing the sensory neuron to spike in proportion to the hair deflection [3]. **C**. Example time course of the joint angle for the *α* joint of the right anterior leg (R1). **D**. Proposed SNN architecture for distributed proprioception of limb and body posture of the stick insect. The joint angles *α, β* and *γ* for each of the six legs are converted into hair angles that are sensed by one mechanoreceptor per hair (Hair field layer). In the second layer, the velocity and position INs encode the posture and movement of individual joints.

In this work, SNNs are used to accurately encode limb kinematic information based on proprioceptive sensory arrays with phasic-tonic response properties (Fig 1D), as evaluated by experimental data comprising concurrent joint angle time courses from 6 × 3 joint angles from unrestrained walking and climbing stick insects (Fig 1C). As outlined above, this is the first of two companion papers investigating fundamental aspects of distributed proprioceptive encoding. Whereas this first part implements phasic-tonic encoding and the extraction of position and velocity in a spiking network, the second part tests to what extent it is possible to extract higher-order parameters like intra-leg movement primitives as characteristic signatures of particular step cycle phases, or inter-leg information concerning whole-body posture, like body pitch (van der Veen et al., companion paper [23]).

The remainder of this work is structured as follows: The Methods section provides an introduction to the dataset and a detailed explanation of the network architecture, describing the methodology layer by layer. The result section presents our findings for each layer of the network. The discussion section provides a comparative analysis with the relevant scientific literature, explores the strengths and weaknesses of the proposed model, and ends with an outlook on future research.

## Methods

### Dataset

The experimental data used in this study was originally collected to study whole-body kinematics of walking and climbing stick insects, comparing related species with different body morphology [44] and characterizing distinct step classes [39]. Stick insects of the species *Carausius morosus* (de Sinéty, 1901) have six legs with fairly similar morphological structure (Fig 1A). Each leg comprises a short basal coxa, a fused trochantero-femur, a long and thin tibia, and distal tarsus with five tarsomeres. The dataset contains motion capture data on three joint angles per leg. Two of these joint angles correspond to joints monitored by proprioceptive hair fields: Hair fields on the coxa (red arrows labelled Cx) measure protraction-retraction movements of the thorax-coxa joint, denoted by a blue *α* in Fig 1A [40, 41]; The trochanteral hair field (red arrows labelled Tr) measures levation-depression movements of the coxa-trochanter joint, denoted by a green *β* in Fig 1A [43]. The dataset also contains extension-flexion movements about the femur-tibia joint, denoted by a red *γ* in Fig 1A. Throughout this work, these three joints are referred to as the *α, β* and *γ* joints, respectively. The front, middle, and hind legs are labeled as 1, 2, and 3 for the right (R) and left (L) sides. Fig 1A also indicates the location of antennal hair fields at the head-scape and scape-pedicel joints on the antenna (labelled by Sc and Pd for scape and pedicel, respectively). Since the dataset does not comprise antennal joint angles, the current study focuses on leg proprioception only. However, given similar function of all proprioceptive hair fields, the spiking network proposed in this work could potentially be expanded to include antennal kinematics. Generally, we assume that the joint angle encoding mechanism follows the same principle for all mentioned joints and all limbs.

The dataset comprises complete body kinematics of unrestrained walking and climbing stick insects. Nine specimens walked freely on a horizontal walkway measuring 40 mm in width and 490 mm in length. In this work, we focus on trials where the animals encountered a flat surface, whereas the companion paper expands to include climbing trials with two stairs [39]. A marker-based motion capture system (Vicon MX10) was employed, using eight infrared cameras capturing 200 frames per second, to track markers attached to the head, thorax, and all six legs of the insect. The captured marker trajectories were used to reconstruct the time courses of the three mentioned leg joint angles *α, β* and *γ*.

### Spiking neural network (SNN) architecture

A schematic of the proposed SNN architecture is shown in Fig 1D. The temporal evolution of each experimentally obtained joint angle is converted into a set of hair deflection angles, corresponding to the number of hairs in the hair field. Each hair has a unique receptive field that is arranged in sequence with receptive fields of adjacent hairs. The combination of all receptive fields results in sensitivity for the entire working range of the joint. In analogy to real proprioceptive hair fields (Fig 1B), each hair deflection is converted into an electric current that is integrated by a single mechanosensory neuron per hair. The mechanosensory neuron consists of an AdEx model [35] and provides a nonlinear phasic-tonic response to a step increase in stimulus magnitude (here: joint angle). This is due to an adaptation mechanism that modulates the relationship between the input current and the output spike rate. The resulting spike trains of all sensory neurons per hair field converge on a set of four first-order INs per joint: two position and two velocity INs. One position IN encodes negative joint displacement relative to rest (denoted as *pos*_−_), while the other encodes positive joint deflection relative to rest (*pos*_+_). Similarly, the velocity INs encode joint movement into either the positive (low joint angle → high joint angle) or negative (high joint angle → low joint angle) direction, denoted as *vel*_+_ and *vel*_−_, respectively. For simplicity, each leg of our model has three joints, each with its own hair plate containing two hair fields with the same proprioceptive encoding mechanism, assuming that key features such as receptive field size, linear range fractionation with equal sensitivity, and phasic-tonic response time courses are the same at all 18 leg joints. Accordingly, the model uses 3 sets of two hair field implementations per leg, totalling 3 × 2 × *N*_h_ mechanosensory neurons that converge onto 3 × 2 = 6 position INs and 3 × 2 = 6 velocity INs. Here, *N*_h_ represents the number of hairs in a hair field. Both IN types are modelled by a simplified LIF model, providing a linear relation between an input current and output spike rate.

The proposed architecture was implemented in Python version 3.9. The simulations were conducted on a system equipped with 16 GB of RAM and an AMD Ryzen 5600x processor. The differential equations that describe the behaviour of spiking neurons and synapses were solved over time using the backward difference method and a time step of *dt* = 0.25 ms.

### Layer one: Hair plate

#### Hair field

It is important to note that while, proprioceptive hair fields are present at the *α* and *β* joints of each stick insect leg, they are absent at the *γ* joint. Despite this fact, hair field models were applied to all *γ* joints as well because mechanosensory neurons of the femoral chordotonal organ, i.e., the sensory organ encoding the *γ* joint angle, share important encoding features with mechanosensory neurons of proprioceptive hair fields, such as sensitivity to joint angle, joint angle velocity [45], and range fractionation [19, 36].

In each hair field, the time course of the respective joint angle is transformed into a set of hair deflection angles, corresponding to the number of hairs on a hair field, *N*_h_. Each hair features a distinctive receptive field arranged in series with its neighboring hairs. The receptive field of each hair is defined as the range of joint angles within which the sensillum is sensitive to changes in deflection. For simplicity, the collective contribution of all hairs spans the entire possible range of the joint angle. Moreover, the receptive field size and the spacing between hairs are uniform for all hairs within a given hair field, deviating from the variation observed in biological hair fields [26]. It is also assumed that there is a degree of overlap between receptive fields. This accounts for the fact that many hair fields are not arranged in regular hair rows but form patches (or plates) of hairs with overlapping receptive fields. Generally, it is unlikely that full deflection of one hair coincides perfectly with the onset of deflection of the next hair. Finally, it is assumed that hair deflection is linearly proportional to the joint angle and the range is bound to [0°, 90°]. If the joint angle falls below or exceeds the proprioceptor receptive field, the hair angle is either not deflected at all (0°) or fully deflected (90°), respectively. This results in the following relation between joint angle (*θ*) and hair angles (*ϕ*_*ij*_) for hair *i* and hair field *j*:

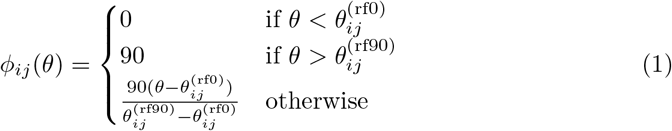

where 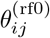 and 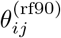 are the lower and upper receptive field edges, respectively, defined as follows:

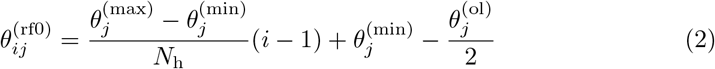

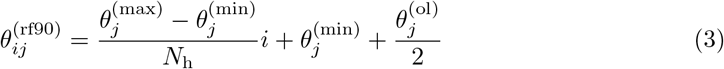

where 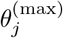 and 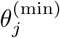 represent the maximum and minimum of the the joint angle sensitivity for hair row *j*, respectively. In the proposed network, these parameters are set as the maximum and minimum joint angles attained by the corresponding joint. Therefore they are not equal for all hair fields. The receptive fields of the outer hairs are manually set to 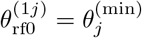 and 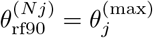. The overlap between two adjacent receptive fields is defined by the parameter 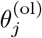 and is equal for all hair fields. Similar to *N*_h_, which is also consistent across all hair fields.

#### Bi-directional hair fields

Hair fields such as the antennal [13] or coxal [41] hair fields of stick insects are often arranged in opposing pairs. Accordingly, our model divides proprioceptive hair fields at each joint into two complementary and opposing subgroups. The parts of these bi-directional hair fields (Fig 2) have their working-range either in the upper or lower half of the joint angle working-range. The halfway point of the joint working-range is called the ‘resting angle’ or ‘neutral angle’, at which no hair is fully deflected. The two hair fields deflect due to increasing and decreasing joint angle relative to the resting angle, respectively. Hair fields with opposing deflection sensitivity require a modification in Eq (1):

**Fig 2.**
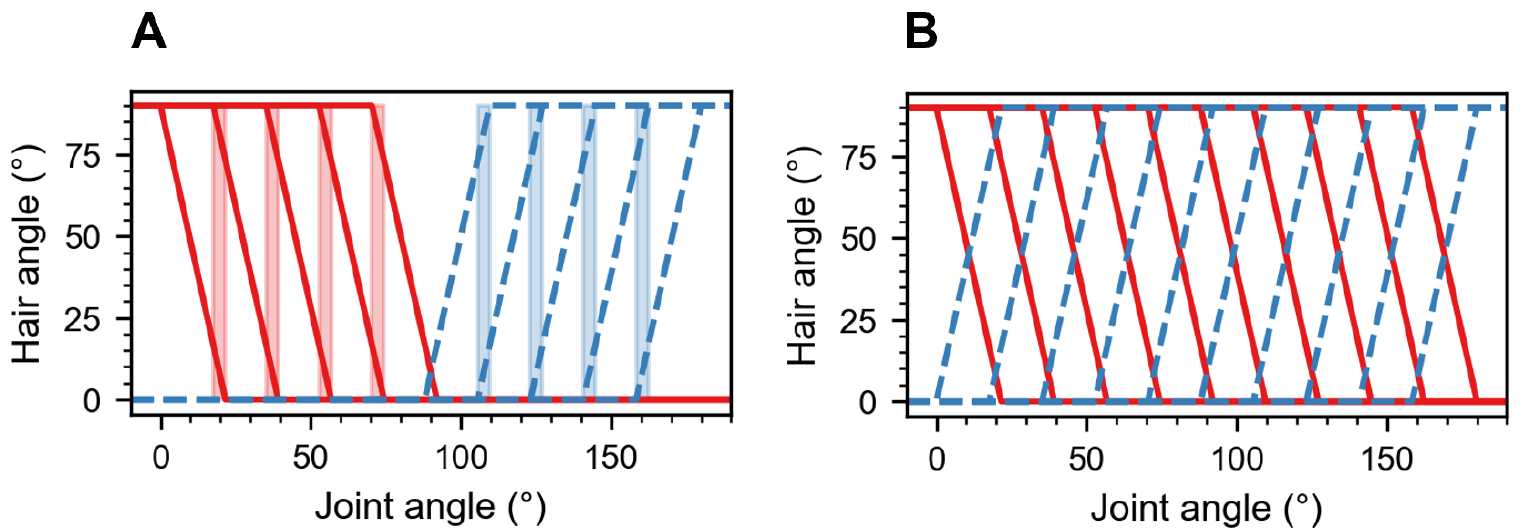
Bi-directional hair field and extended bi-directional hair field. **A**. The bi-directional hair field models pairs of hair rows on opposite sides of the joint and working-ranges below (red) and above (blue) the neutral angle 90°. In the idealised hair row, hair angle is a function of joint angle with parameters 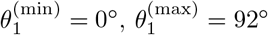, 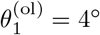, and *N*_h_ = 5 (solid red lines), and 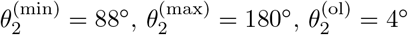, and *N*_h_ = 5 (dotted blue lines). The positively and negatively oriented hair angles are calculated using Eq (1) and Eq (4), respectively. The resting angle is at 90°. **B**. Extended bi-directional hair plate: This hypothetical scenario is modified from panel A as follows: *N*_h_ = 10, 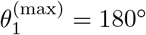 and 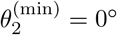. Overlap ranges have been omitted for clarity.

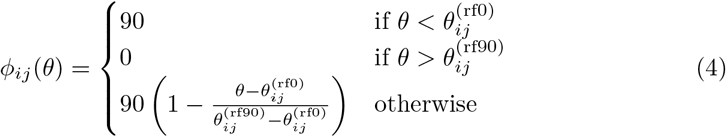

Moreover, the calculations for 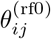 (Eq (2)) and 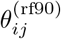 (Eq (3)) are interchanged.

Fig 2A illustrates multiple hair angles in relation to the joint angle for a hypothetical scenario. The calculations for positively and negatively oriented hairs are determined using Eqs (1) and (4), respectively. The positively oriented hair field is sensitive from 88° → 180° while the negatively oriented hair field is sensitive from 92° → 0°.

Incorporating an overlap into the hair plate was intended to maintain low but non-zero spike rates at the resting position for the hair field. The overlap between the hair fields (*θ*^(olhf)^) at the resting position (90° in the hypothetical scenario) is defined as follows:

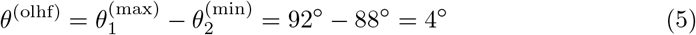

In the proposed network, the parameters *θ*^(olhf)^ and 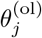 are set to the same value.

#### Adaptive Exponential Integrate-and-Fire (AdEx) model

A deflected hair acts as a lever arm and applies force to the tip of the sensory neuron’s dendrites, opening mechanotransduction channels and generating a current [46]. To reflect this in the model, the hair angles calculated in Eqs (1) and (4) were multiplied by 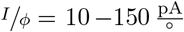 yielding currents in the nA range. This current flows into a single sensory neuron, converting the input current into a spike train [3]. To model the mechanosensory neuron dynamics of insect hair fields, we used empirical data on the dynamics of antennal proprioceptive hair field afferents characterized by Okada and Toh (2001) [38], illustrated in Figs 3A,B. These experiments were carried out on the antenna of the American cockroach *Periplaneta americana*. Therefore, the mechanosensory mechanisms can be assumed to be similar between species and location, although particular parameters may need to be adapted to reflect differences in biology. Multiple neuron models were found suitable to replicate these non-linear spiking dynamics [47]. While carefully considering biological accuracy, implementation costs, and potential spiking dynamics, the Adaptive Exponential Integrate-and-Fire (AdEx) model was chosen as the mechanosensory neuron model [35]. The AdEx model dynamics are governed by two ordinary differential equations (ODEs):

**Fig 3.**
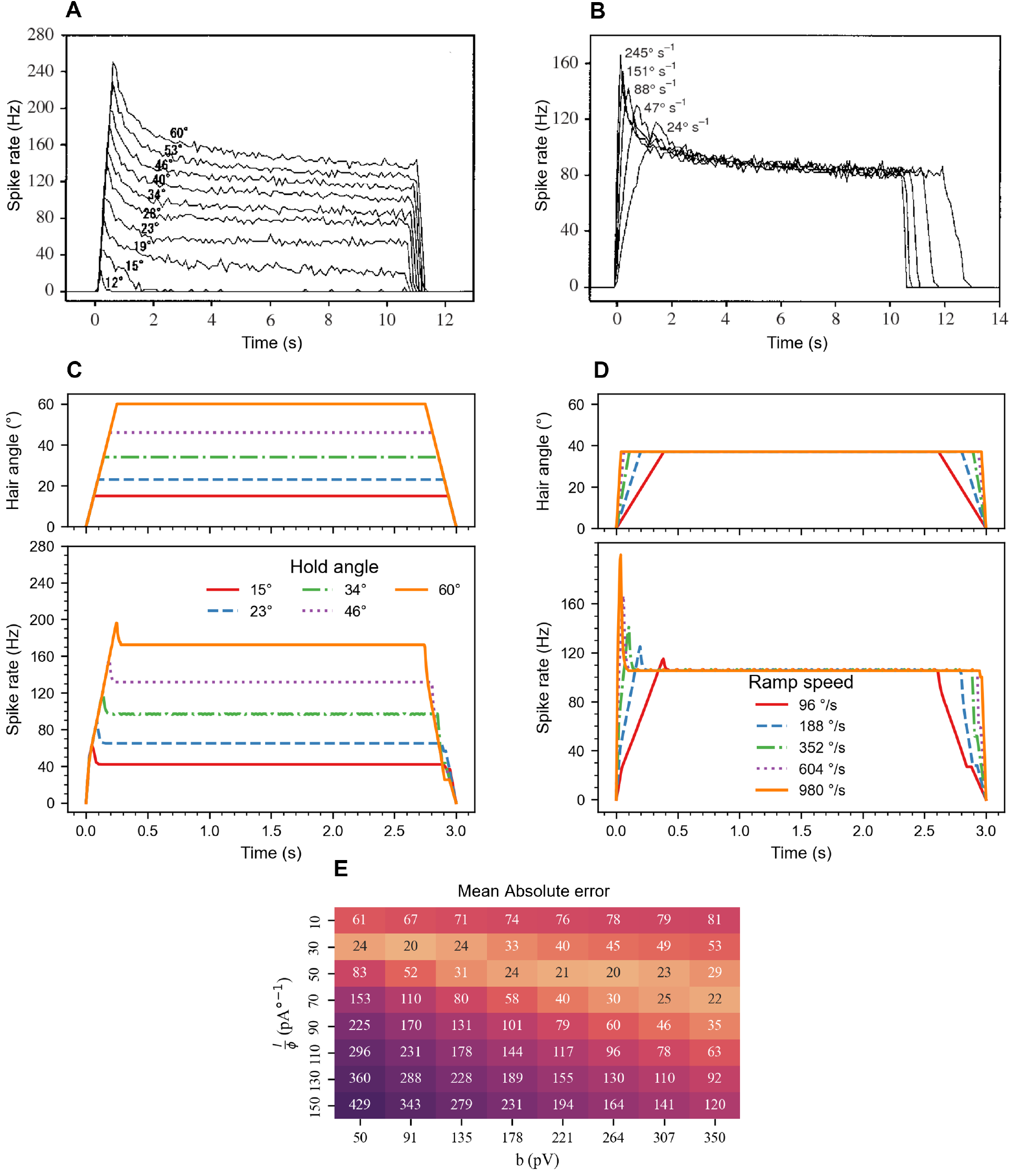
Phasic-tonic spike response of the real and modeled mechanosensory neuron. **A**. Spike response of mechanosensory afferents from the lateral scapal hair plate in the American cockroach [38]. The plot shows the spike rate over time for a tactile hair deflected by a ramp-and-hold function with a constant velocity of 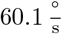 from 0° to various end angles. **B**. Response to ramp-and-hold deflection of a tactile hair with a constant hold angle of 37°, but varying angular velocities. **C**. Stimulus protocol (top) and corresponding spike responses of the AdEx model (bottom). The stimulus protocol is the same as for A, except with four-fold ramp velocity and shorter hold time. **D**. Stimulus protocol (top) and corresponding spike responses of the AdEx model (bottom). The stimulus protocol is the same as for B, but with four-fold ramp velocities and shorter hold time. **E**. Heatmap illustrating the mean absolute error (MAE) between significant spike rates (peak and steady-state) of the modeled and experimental sensory neuron for a parameter sweep across *I* / ϕ and *b*. The MAE was computed according to Eq (12).

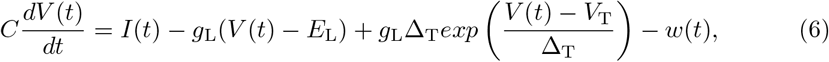

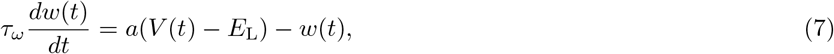

where *V* (*t*) represents the membrane potential, *C* denotes the capacitance, *I* the input current, *g*_L_ the leak conductance, *E*_L_ the leak reversal potential, Δ_T_ the slope factor, *V*_T_ the threshold voltage, *w*(*t*) the adaptation variable, *a* the adaptation coupling factor, and *τ*_*w*_ the adaptation time constant [35].

In a real neuron, an action potential (spike event) occurs due to depolarization. In the AdEx model, when *Vs* > *V*_t_, a spike is recorded, and the timestep is noted as *t*^*f*^. At this timestep, *V* (*t*) is reset to *E*_L_ and *ω*(*t*) increases by a constant *b*.

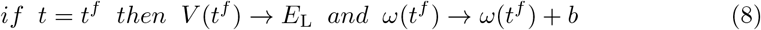

The second term in Eq (6) on the right-hand side (RHS) represents the passive membrane function, implemented as a leakage mechanism that allows the membrane voltage to return to *E*_L_, in the absence of an applied current. In a biological neuron, this leakage term represents the random diffusion of ions across the membrane. The third term allows for the active (or voltage-dependent) membrane properties, whereby the membrane voltage triggers an exponential spike if it exceeds the threshold voltage, resulting in a transient, positive overshoot of the membrane potential. The fourth term on the RHS corresponds to the adaptation term, which modulates the active membrane properties by means of a current flowing out of the membrane. Consequently, an increase in *w*(*t*) results in a decrease in the sensitivity of *V* to a stimulus. The adaptation variable increases when the membrane voltage exceeds its resting state (Eq (7)) or after a spike event (Eq (8)). The second term on the RHS of Eq (7) allows the adaptation variable to converge to zero in the absence of spikes, thereby resetting the adaptation process. Therefore, adaptation reduces the neuron’s sensitivity to a stimulus due to a prolonged or repeated stimulus. For reference, the response of the AdEx model to a constant current stimulus is shown in S1 Fig.

### Layer two: Position interneurons (INs)

The overall activity of the hair plate could be decoded as the temporal evolution of joint angles by an IN that integrates spikes from all proprioceptors in a hair field. Due to the binary nature of the hair field (Fig 2A), two position neurons were linked to the sensory neurons of negatively and positively oriented hair fields, respectively. With this arrangement, only one position neuron becomes active depending on whether the joint is within the negative or positive working range relative to its resting position, similar to the position encoding INs identified by Ache and Dürr (2013) [19].

#### Leaky Integrate-and-Fire (LIF) model

Due to its integration capabilities, simplicity, and small parameters set, the LIF model was chosen as the position IN model [47]. In this model, a pre-synaptic spike that occurs at time *t*^pre^ increases *V* (*t*) by a synaptic weight *ω* [48]:

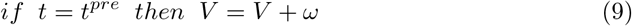

The dynamics of the LIF are as follows:

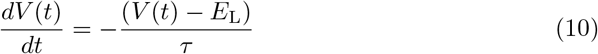

where *τ* is the decay time constant. If the voltage *V* (*t*) exceeds the threshold voltage *V*_T_, a spike time was recorded as *t*^*f*^ and the voltage was reset to *E*_L_ at this timestep:

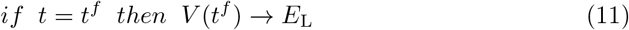

The only tunable parameters are *τ* and *ω*. For reference, the response of the LIF model to a constant frequency spike train stimulus is provided in S2 Fig.

### Layer two: Velocity interneurons (INs)

First-order mechanosensory INs can also encode joint movements. For example, the spike rate of a movement-sensitive IN of the antennal mechanosensory system increases linearly with joint velocity but remains inactive when the joint is stationary [37]. To replicate this kind of velocity encoding, velocity INs are modeled to spike whenever the joint angle transitions from the receptive field of one hair to the next. To accomplish this, we could leverage the phasic response of the sensory neurons.

#### Modified hair field distribution

Similar to the position INs, the movement layer features two velocity INs per joint. The INs fire during an increase (*vel*_+_) or decrease (*vel*_−_) of joint angle, respectively. However, to encode velocity across the entire working range, both neurons need to have sensitivity throughout the complete joint range of motion. Owing to the directional selectivity of each hair’s response, this is not achievable with the opposing arrangement of the simple bi-directional hair field. Consequently, the scenario shown in Fig 2A was extended by including 5 additional hairs and expanding the joint range for both hair fields: *N*_h_ = 10 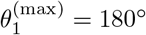, and 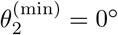. This modification is hypothetical in that it could either account for a response to ‘release of deflection’, or for a fraction of hairs being fully deflected at the rest position of the joint. Both of these properties would introduce bi-directional selectivity for the entire joint angle range (Fig 2B), omitting overlap surfaces for clarity. Note that the supplementary hairs of the extended bi-directional hair field were not linked to the position IN.

#### Neuron model for a high-pass filter

The mechanosensory neuron of each tactile hair has a phasic response at the moment when the hair reaches its maximum deflection, resulting in the spike rate peak shown in Figs 3A,B. For fully deflected hairs (*ϕ* = 90°), the steady-state spike rate of the sensory neuron (*f*_ss_) is constant for all hairs at all times. In order to exploit the phasic behaviour of the sensory neuron, a single high-pass filter is integrated in series with the sensory neuron, whose cut-off frequency (*f*_*c*_) is just above *f*_ss_. Consequently, only the phasic response of the sensory neuron can trigger spikes from the high-pass filter. And thus, the neuron only fires when the joint angle exceeds the receptive field of the tactile hair and the hair reaches its maximum deflection. The joint velocity can then be extracted from the collective spike rate of the high-pass filters.

The LIF model, governed by Eqs (9), (10), and (11), can operate as a high-pass filter [49]. Due to the leaky term in Eq 10, the LIF only spikes if a specific input frequency is reached (*f*_*c*_). The cut-off frequency, *f*_c_, depends on both *ω* and *τ*. By setting *τ* to a specific value and manually adjusting *ω, f*_c_ can be set slightly above *f*_ss_. Filtered spikes from a hair field converge onto a single LIF neuron, which generates a spike in response to each input spike. This IN integrates the spike rates into a unified output, representing the velocity. A LIF generates a spike in response to a single input spike if *ω* > *V*_T_ − *E*_L_.

## Results

### Layer one: Hair field layer

Within the hair field layer, each joint was associated with two distinct hair fields, each containing a total of *N*_h_ individual hairs. Within a hair field, each hair was sensitive to joint deflection in its own receptive field. Within this receptive field, the hair deflection angle was translated into an electrical current and transmitted to a single mechanosensory neuron designated to that hair. Subsequently, the mechanosensory neuron transformed a increment change in current into a phasic-tonic spiking response. This process mirrored the mechanosensitive dendrites found in actual tactile hair sensory neurons (Fig 1B) [3].

#### Replicating sensory spiking dynamics

The AdEx neuron was used to model the electrophysiological dynamics of tactile hair sensory neurons observed by Okada and Toh (2001) [38], as depicted in Figs 3A,B. This was achieved by replicating their experimental procedure and adjusting model parameters to match the observed spiking dynamics. To achieve similar adaptation dynamics in the model, a relatively high *τ*_*ω*_ (approximately 600 ms) was required. However, prolonged adaptation adversely affected accuracy in the following layer, as adaptation persisted beyond the desired time frame. Consequently, *τ*_*ω*_ was manually set to 50 ms, and the corresponding angular velocities of hair deflection were quadrupled by dividing the total stimulus duration by four. This adjustment allowed us to replicate the general dynamics of the phasic-tonic response, albeit with different adaptation speeds. The modified procedures from the study of Okada and Toh (2001) [38] were:

1. A ramp-and-hold function with linear increase of hair deflection at different speeds, while keeping the hold angle constant. Hairs were deflected from 0° to 37° at five velocities. These were 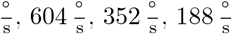 or 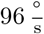
2. A ramp-and-hold function linear increase of hair deflection at constant speed, but differing hold angles. Deflection velocity was 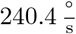 The five hold angles used were: 60°, 46°, 34°, 23°, or 15°.

A grid search was employed to vary the values of *I / ϕ* and *b* within the ranges of 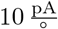to 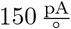 and 50 pV to 350 pV respectively, with a total of 8 steps. The parameter *I / ϕ* influences the current entering the neuron, thereby affecting the response strength. The parameter *b* modulates the degree of adaptation, impacting the relative strength of the phasic peak. Together, these parameters regulate both the phasic peak and steady-state frequencies, which are the key response metrics of the model. Several other parameters can be modified to achieve the desired spiking response (see S3 Fig, S4 Fig, S5 Fig). The error metric mean absolute error (MAE) was computed to assess the deviation of the model response from the electrophysiological dynamics observed in the experimental study for critical spike rates (maximum and steady state), with:

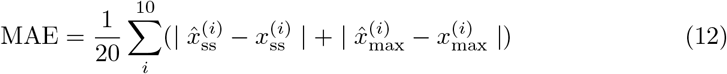

for the *i*th ramp function trial. 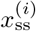 refers to the spike rate at the steady state, 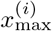 represents the maximum spike rate, and the notation 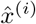 indicates values for the AdEx model while *x*^(*i*)^ indicates values extracted from Figs 3A,B.

#### Optimization and performance

Before optimizing *I / ϕ* and *b*, the initial AdEx parameters were adopted from Naud et al. (2008) [50]. In order to account for the high sensitivity of the mechanosensory neuron to small hair deflections, *g*_L_ was reduced from 12 nS to 2 nS. This decreased the rheobase current, enabling the neuron to detect weak input currents. Additionally, the time constant *τ*_*ω*_ was reduced to 50 ms, as explained in the previous paragraph. Fig 3E shows the MAE (computed using Eq (12)) in each iteration of a parameter sweep over *b* and *I / ϕ*. A minimum error of 20 is observed for several results, from which the values 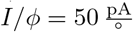 and *b* = 264 pV were randomly selected. The optimal parameters for the AdEx model are listed in Table 1.

**Table 1.** Model type and parameter values for the sensory neuron, position INs and velocity INs after optimization.

| IN | Type | $C$ | $g_L$ | $E_L$ | $\Delta_T$ | $V_T$ | $a$ | $\tau_w$ | $b$ | $\tau$ | $\omega$ |
| --- | --- | --- | --- | --- | --- | --- | --- | --- | --- | --- | --- |
| Sensory | AdEx | 200 pF | 2 nS | -70 mV | 2 mV | -50 mV | 2 nS | 50 ms | 264 pV | - | - |
| Position | LIF | - | - | -70 mV | - | -50 mV | - | - | - | 120 ms | 1 mV |
| Velocity | LIF | - | - | -70 mV | - | -50 mV | - | - | - | 5 ms | 10.8 mV |

The results of both hair deflection experiments are illustrated in Figs 3A,C (varying the hold angle) and 3B,D (varying angular velocity of the ramp) using the optimal parameter set. The dynamics can be directly compared to the experimental data on hair field afferents of the lateral scapal hair plate of the American cockroach (Figs 3A,B) [38]. The stimulus protocols employed for the experimental and model data were comparable, although the former incorporated four-fold angular velocities and a shorter hold time. Despite these adjustments, the modelled sensory neuron exhibited phasic-tonic dynamics similar to that observed in the experimental data. During the ramp phase of the stimulus, the spike rate showed a linear increase, reaching a peak as the ramp reaches the constant hold angle. Subsequent to the peak, the spike rate converged to a steady state. As the stimulus was ramped back to zero, the spike rate showed a linear decrease until it reached zero. In contrast to the considerable noise present in the experimental data, the model is deterministic and, therefore, devoid of noise. Furthermore, the model reached a steady state within 100 ms, whereas the experimental data displayed a continuous decay throughout the entire hold phase.

In trials with varying hold angles (Fig 3A,C) the transient peaks reached approximately 110 % of the steady-state conditions in the model. In contrast, the peaks in the experimental data were approximately 150 % of the steady-state spike rate. Moreover, in the experimental data, the low-end deflection angles (e.g., *θ* = 15°) only produced a response during the onset of the ramp function, not during the steady state. This behavior was not observed in the simulated model, where low deflection angles induced a steady-state spike response. Except for very small hold angles, the steady-state spike frequencies of the model aligned closely with experimental results.

Trials with varying ramp velocities (Figs 3B,D) exhibited a higher degree of similarity. The steady-state spike rate was slightly higher than the noise range of the experimental data. Additionally, the peaks were in a slightly wider range: 108 Hz - 199 Hz for the model compared to 118 Hz - 166 Hz for the experiment.

### Layer two: Position interneurons (INs)

This section commences with an evaluation of the characteristics of the bi-directional hair field, which is followed by an analysis of the performance of the position INs. Two position INs were integrated in each joint, with the function of integrating all spikes originating from a given hair field into a position-dependent spike train. These spike trains were able to encode the varying joint angles over time, through their spike rate.

#### Bi-directional hair field

In each hair plate, 100 hairs were arranged to form two hair fields, each comprising 50 hairs (*N*_h_ = 50). For each joint, the bounding joint angles of the bi-directional hair field, 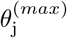 and 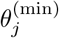, were set as the maximum and minimum values of the joint’s working-range. The overlap was defined as follows: *θ*^(olhf)^ = *θ*^(ol)^ = 0.1°. As was clarified in the methods section, half of the hairs in a hair field were connected to position INs. Accordingly, Fig 4A included only hairs with synaptic connections to position INs. The time-varying plot displays a 5-second time course for the *α* joint angle of the right front leg, R1. The dotted line represents the resting angle, situated in the middle of the joint range. Red and blue dot sequences form a hair field raster plot, where each dot denotes a spike event for the *i*th hair, respectively. Due to the design of the hair field, proprioceptive hairs numbered 1 to 25 are sensitive to joint angles in the negative domain, while hairs 51 to 75 are sensitive to joint angles in the positive domain. Due to the large number of spikes, dots representing individual spike events coalesce into a continuous line.

**Fig 4.**
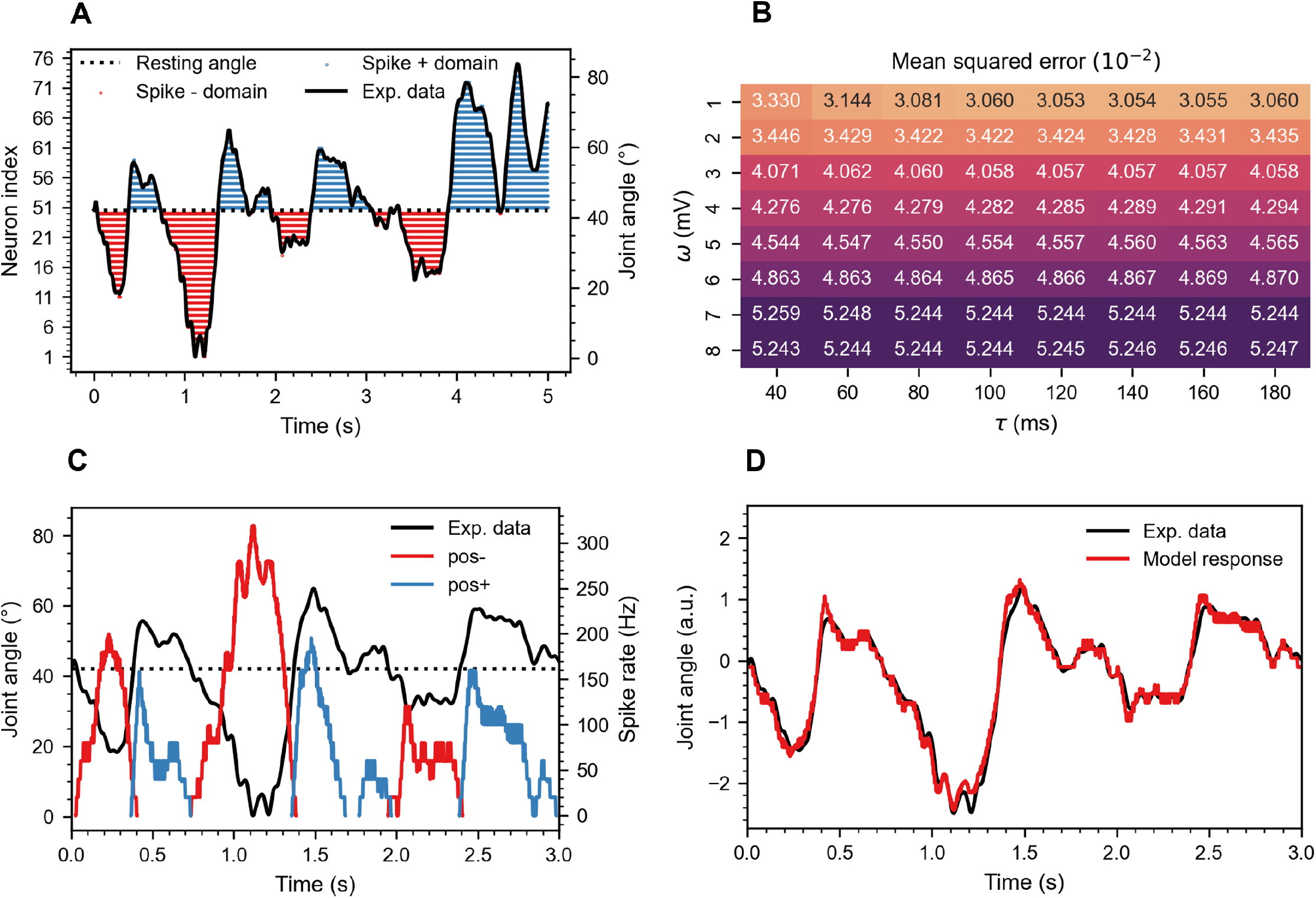
Position IN results. **A**. Raster plot displaying the response of a hair plate (*N*_h_ = 50) to a 5-second movement sequence of the *α* joint (thorax-coxa) of the right front leg (R1). Spike events from proprioceptive hairs in the anterior (blue dots) and posterior (red dots) parts of the joint’s working range encode increasingly protracted or retracted postures, respectively. The high spike density results in the dots merging into a continuous line. **B**. Heat map depicting the mean squared error (MSE) between experimentally obtained joint angles and model predictions across a parameter sweep of *τ* and *ω*. The results are averaged over 78 trials and 18 joints, totaling 1404 joint angle time courses. **C**. Spike rate time courses of the *pos*_+_ and *pos*_−_ INs in response to the same joint angle movement as shown in panel A. **D**. Combined and z-normalized spike rates of the *pos*_+_ and *pos*_−_ INs from panels A and C closely follow the joint angle time course. The model response and experimental data are plotted in arbitrary units due to z-normalization.

Fig 4A highlights the bi-directional sensitivity of the hair field, with mirrored encoding observed in the positive and negative domains. In the positive domain, increasing joint angles prompted additional neurons to activate, whereas in the negative domain, the opposite was true. At the resting angle, only the initial sensory neurons (25 and 26) activated due to the pre-defined overlap. Ultimately, the close alignment between joint angle and hair field activity was evident.

#### Optimization and performance

To assess the performance of the position INs, we interpolated the experimental joint angles to match the timestep of the model (*dt* = 5 ms → 0.25 ms). Subsequently, we subtracted the spike rate of the *pos*_−_ IN from the *pos*_+_ IN and then z-normalized the resulting time series 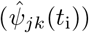. For direct comparison, we z-normalized the corresponding joint angle time course (*ψ*_*jk*_(*t*_i_)), as illustrated in Fig 4D. The squared difference between these z-normalized time courses was then averaged over each discrete time step *i*, joint angle *j* and trial *k*, yielding the MSE for one joint angle time course:

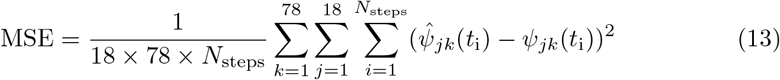

The MSE, averaged over 78 trials and 12 joint angles, was computed at each iteration of a parameter sweep for *τ* and *ω*, and the results were visualized in a heat map (Fig 4B). The optimal parameters, *τ* = 120 ms and *ω* = 1 mV yielded an MSE of 0.030 55. A decrease in MSE was observed while lowering *ω*. The synaptic weight was not reduced further because the maximum observed spike rate (Fig 4C, *f*_*max*_ = 322 Hz) was already low. Further reduction would have introduced more stair stepping. The time constant *τ* influenced the MSE little at *ω* values of 2-10 mV. While for *ω* = 1 mV, low *τ* values increased the MSE. At these values, the spike rate was too low to allow for detailed encoding by spike rate modulation, due to pronounced stair stepping. The complete optimal parameter set of the position IN is listed in Table 1. Fig 4C illustrates a 3-second joint angle time course and spike rates for the *α* joint of R1. The *pos*_+_ and *pos*_−_ neurons fired proportionally to the joint angle relative to the resting angle. Spike rates fluctuated between 0 and 322 Hz and followed the joint angle time courses (though inverted in case of the hairs in the posterior working-range). Note that the degree to which fine details of the joint angle time course were lost in spike rate modulation depended, apart from the model parameters mentioned above, on the number of proprioceptive sensory neurons per hair field. If spike rates in the kHz range are required for further processing, as is the case for the subsequent layer in the companion paper (van der Veen et al., companion paper [23]), *ω* can be increased (e.g., *ω* = 25 mV for the companion paper). To visually assess joint angle encoding, the response of *pos*_−_ was subtracted from the response of *pos*_+_ and then z-normalized. This time course is plotted alongside a z-normalized joint angle time course in Fig 4D. Visually, the spike rate closely resembled the joint angle time course. However, the model response ‘overshot’ the joint angle time course at moments of sudden changes in joint angle (e.g., *t* = 0.5 s, 1.5 s, 2.5 s). Additionally, there was some stair stepping in the spike rate.

To test whether position IN performance was the same irrespective of leg or joint types, a two-way Analysis of Variance (ANOVA) [51] was conducted to examine the effect of the factors ‘joint type’ (*α, β* and *γ*) and ‘leg type’ (front, middle, and hind) on the MSE. The results showed no significant main effect of factor ‘leg type’ (F(2, 69) = 2.52, p = 0.084). However, there was a significant main effect of factor ‘joint type’ (F(2, 69) = 111.62, p = 9.62 × 10^−32^), indicating that position encoding varied significantly among joints. A post hoc comparison using the Tukey’s honestly significant difference (HSD) test [52] indicated that the MSE of the *β* joint was significantly different than the MSE of the *α* and *γ* joints. Table 2 lists the specific MSE values for the three leg types and three joint types. The table reflects that position encoding was statistically indistinguishable between leg types, but significantly worse for the *β* joint compared to the *α* and *γ* joints. Additionally, a significant interaction between leg types and joint types was observed (F(2, 69) = 11.11, p = 4.90 × 10^−8^). A post hoc comparison using the Tukey’s HSD test indicated that for the *α* joint, the front legs performed significantly worse, and for the *β* joint, the middle legs performed significantly worse.

**Table 2.** MSE and standard deviation (SD) for the position INs, grouped by factors legs and joints of the two-way ANOVA.

| Leg | MSE | SD | Joint | MSE | SD |
| --- | --- | --- | --- | --- | --- |
| front | 0.0317 | 0.0071 | $\alpha$ | 0.0266 | 0.0046 |
| middle | 0.0310 | 0.0105 | $\beta$ | 0.0389 | 0.0075 |
| hind | 0.0296 | 0.0060 | $\gamma$ | 0.0268 | 0.0047 |

### Layer two: Velocity interneurons (INs)

The objective of this section is to test how a velocity-sensitive first-order IN could encode joint angle velocity independent of joint position, despite its presynaptic input elements being position-sensitive. As with position INs, the instantaneous joint angle velocity could be encoded through spike rate modulation. With regard to the encoding properties of the mechanosensory neurons, it could be observed that joint movement was signalled by transition of joint angle from one hair’s receptive field to the next. In order to encode such transition events in a systematic manner, we decided that the phasic response of the sensory neurons should be leveraged. The phasic response component of a hair field afferent was driven by the initial deflection of the hair, before the adaptive current of the AdEx model built up. It was possible to integrate several phasic response components within an entire hair field in a first-order IN by emphasizing subsequent joint angle transitions from one receptive field to the next.

#### Optimization and performance

First, parameters were optimized to set the high-pass filter properties of the velocity IN such that the cut-off frequency *f*_c_ was just above the steady state frequency *f*_ss_. Since *f*_c_ depends on both *τ* and *ω, τ* was arbitrarily set to 5 ms, and the working-range for *ω* was established through trial and error to be 9.0-10.8 mV.

Fig 5A illustrates a linear relationship between the spike rate of a velocity IN and joint angular velocity. The linear velocity dependency was very much like that velocity-sensitive descending INs iONv and cONv of the stick insect antennal mechanosensory pathway (Fig 4 in [37]), except for the lacking resting activity. For low *ω* (9 mV) or *N*_h_ (25) values, angular velocity was not or weakly encoded at 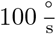 and below. Increasing *ω* and *N*_h_ resulted in a higher overall spike rate, encoding of low velocities and stronger linearity (higher *r*^2^ value). Considering most accurate, linear encoding of velocity, *N*_h_ = 50 and *ω* = 10.8 mV were selected as the optimal parameter set. For these parameters, the spike rate increased with a slope of 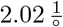 All model parameters of the optimal velocity INs are listed in Table 1.

**Fig 5.**
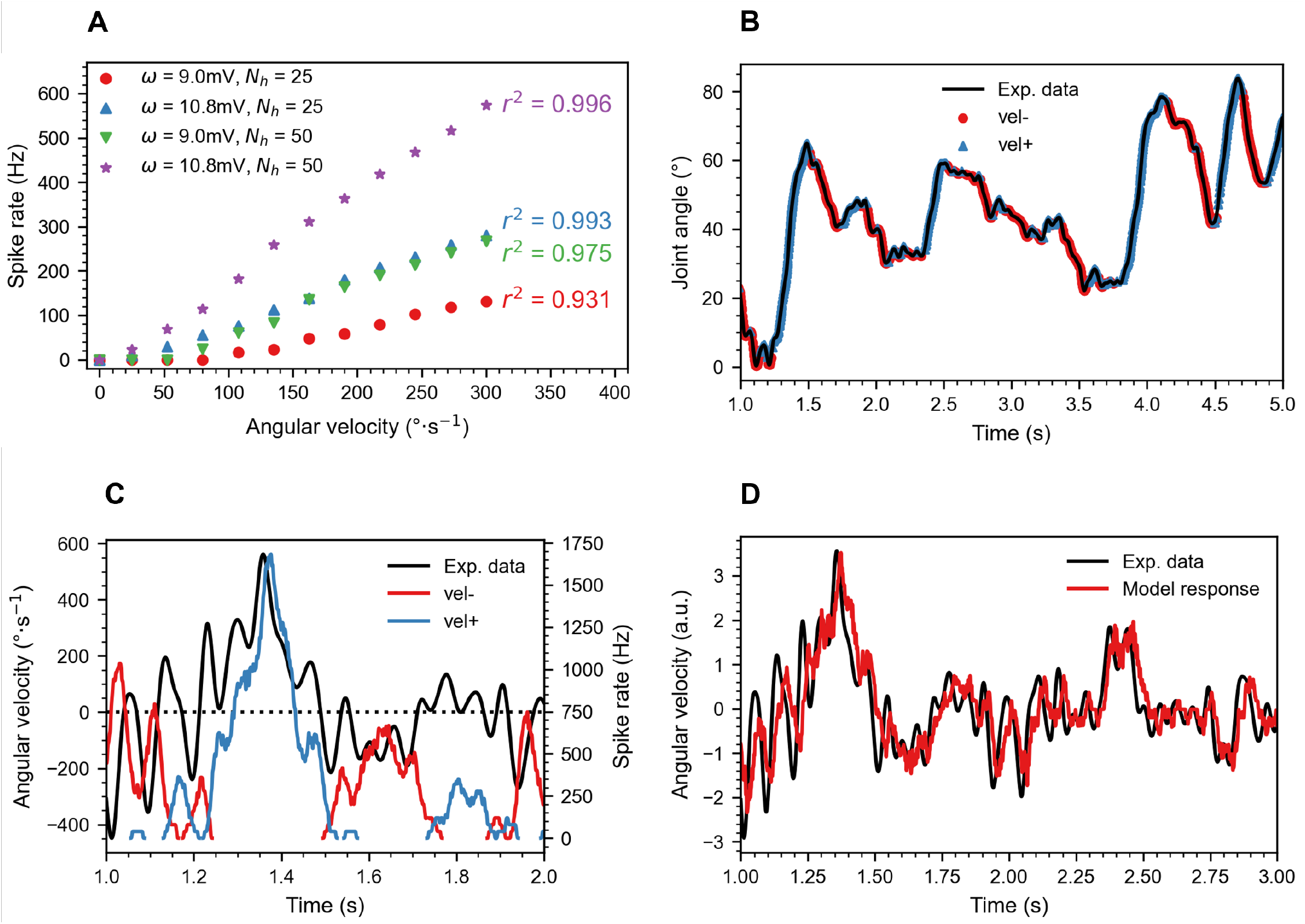
Velocity IN results. **A**. For velocity INs, the output spike rate increases linearly with angular velocity. *ω* and *N*_h_ were varied to test their impact on the spike rate. The four variants shown here differ in threshold velocity for reliable encoding and the slope of the linear relationship. The *r*^2^ value of a linear regression fit is plotted for each variant. **B**. Individual spike events superimposed on the joint angle time course reveal consistent encoding of positive slope by *vel*_+_ INs (blue) and of negative slope by *vel*_−_ INs (red). **C**. The spike rate of the *vel*_−_ and *vel*_+_ INs (red and blue, respectively) in response to a concurrent change in joint angle velocity (black). **D**. Combined and z-normalized spike rate of both the *vel*_+_ and *vel*_−_ INs in response to a representative experimental angular velocity record. Both the model response and experimental data are plotted in arbitrary units due to z-normalization. A persistent time lag of approximately 0.025 s is observed in the model response.

Fig 5B plots individual spikes for the *vel*_−_ and *vel*_+_ INs, overlaid on the joint angle time course for the *α* joint of the right front leg, R1. Evidently, the *vel*_−_ and *vel*_+_ neurons spike during negative and positive joint movement, respectively. The figure illustrates a delayed offset after a sudden changes in joint angle (peak or trough). During these changes, the current in the mechanosensory neuron can decrease but remain high. If adaptation has not fully occurred, the sensory neuron spike rate can still exceed *f*_c_. Consequently, a velocity IN may continue to fire for a short duration even after the joint angle has changed direction, leading to the observed delayed offset and reduced accuracy. Additionally, since velocity IN spikes only occur near the edges of the receptive field, there is no sensitivity to movements within the receptive field, further lowering accuracy.

Fig 5C plots the angular velocity over time in conjunction with the spike rate for the *vel*_+_ and *vel*_−_ INs. Fig 5D plots the combined and z-normalized spike rates of the *vel*_+_ and *vel*_−_ INs. The model response lags behind the experimental data by Δ*t* ≈ 0.025 s. Since the error measures of the position and velocity INs is similar, their values can be directly compared. Overall, velocity INs (MSE = 0.3421) are less accurate than position INs (MSE = 0.030 55). If shifted by 0.025 s, to correct for the lag, the MSE improves to 0.0910, which is still by 0.0605 higher than in position INs.

To further assess the accuracy of the model, the velocity INs were treated as binary classifiers. Spikes of the *vel*_+_ type IN during positive and negative movements (i.e., increasing and decreasing joint angles) were denoted as true positive (TP) and false positive (FP), respectively. Similarly, spikes of the *vel*_−_ type IN during positive and negative movements were labeled as false negative (FN) and true negative (TN), respectively. Since the number of Positive (P) and Negative (N) occurrences was expected to be balanced, a simple accuracy calculation was sufficient:

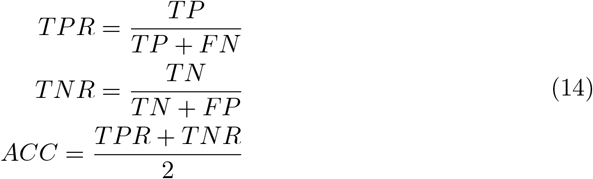

where the true positive rate (TPR) is the sensitivity and the true negative rate (TNR) is the specificity. An accuracy of zero indicates that all spikes occur at incorrect times, while an accuracy of one implies that all spikes occur at the correct time. The averaged results over 18 joint angles and 78 trials are provided in Table 3. The accuracy is high (0.914), and both TPR and TNR are balanced, indicating that both neurons (*vel*_+_ and *vel*_−_) perform similarly accurate.

**Table 3.**
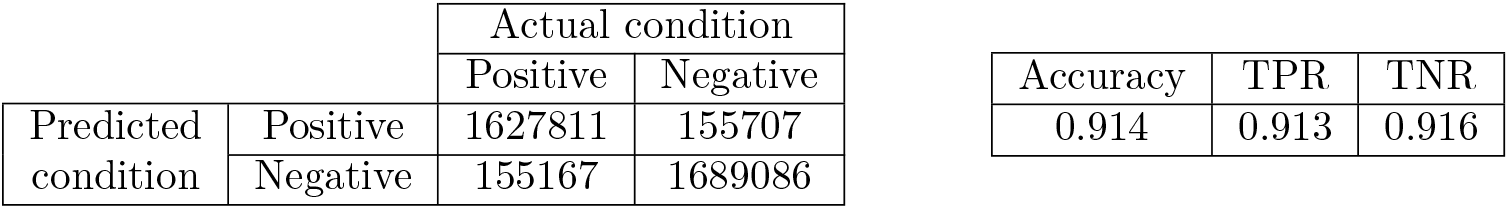
Confusion matrix and accuracy metrics for the velocity INs if regarded as a binary classifier.

## Discussion

Our results demonstrate the use of computationally compact integrate-and-fire neuron models for distributed encoding of joint kinematics. Our focus on proprioceptive hair fields as mechanosensory inputs does not limit the generality of model for distributed proprioception in insects because of computational features that are common to most insect proprioceptors: The first of this concerns the phasic-tonic encoding of the sensory magnitude and associated spike rate adaptation, as captured by the first layer of our model (Figs 2, 3). The second concerns the encoding of position (Fig 4) and velocity (Fig 5) in first-order INs of layer two, despite mixed position-velocity encoding of their presynaptic inputs.

### Layer one: Hair field

Proprioceptive hair fields come in two types, either as two-dimensional patches (hair plates) or linear rows of hairs (hair rows), sometimes with both types occurring on the same limb segment (e.g., [13]). As hair fields are often arranged in pairs at opposite sides of the same limb segment, the sensory array of our hair field layer was designed as a pair of opposing hair rows. Accordingly, the bi-directional hair field model in Fig 2 is a structural template of real hair fields in that the angular working-range of the joint in sampled by two complementary, idealised sensory arrays with equal spacing of hairs.

Structural idealisations of our sensory array concern, for example, the uniform length of tactile hairs, neglecting known length variation [26]. Secondly, a linear relation between hair angle and joint angle was assumed, leading to a linear dependence of tonic spike rate and joint angle, and neglecting non-linearities such as the threshold deflection angle for tonic spike activity (see Fig 3A and below). Finally, our assumption of a linear sensory array (hair row) neglects overlapping receptive fields for multiple hairs that occur for two-dimensional hair plates. Computationally, it is easy to expand our structural template so as to include non-equal spacing and/or multi-hair overlap. On the other hand, there is evidence for tactile hairs being arranged at regular intervals [3, 53].

The phasic-tonic dynamics of a position-dependent response with spike rate adaptation was modelled after experimental data from afferents of the lateral scapal hair plate of the American cockroach (Figs 3A,B) [38]. Other than earlier studies that modelled afferent hair field spikes with a uniform random number generator [34] or Poisson process [54], we employed an integrate-and-fire neuron model. This model family captures the non-linear transformation that occurs between the stimulus (e.g., deflection of the hair) and the neuron’s response, including sub-threshold temporal integration and a spike threshold that is set by a biologically meaningful parameter rather than by mathematical parameters (as in a Poisson process). The phasic-tonic properties associated with spike rate adaptation were incorporated through the AdEx model (Figs 3C, D). In contrast to non-spiking models of non-linear afferent dynamics [33], the AdEx model directly generates spikes rather than capturing only changes in spike rate. Moreover, it accepts raw input of joint angles without undergoing pre-processing via high-pass and low-pass filters, as in [34]. Here, the sole pre-processing step involved is converting joint angles into hair deflection angles, subsequently transforming them into an input current.

The adaptation term in the AdEx model allowed for a combined phasic-tonic response and therefore copied the essential features of the observed experimental spike dynamics, though in a mathematically compact manner. As a trade-off compared to more physiological Hodgkin-Huxley-type models of mechanotransduction (e.g., for spider slit sensilla; [30]) integrate-and-fire models are not suitable for sensitivity analyses addressing the function of particular ion channels.

The spike rate adaptation in the physiological recordings from cockroaches (Figs 3A,B) proved to be relatively slow. Capturing this in the AdEx model required a long time constant (*τ* ≈ 600 ms), which led to significant delays in subsequent layers. After a sudden change in joint angle, the mechanosensory neuron remained suppressed due to adaptation for a relatively long time which, in turn, lead to inconsistent and worse performance in the velocity and position INs. To alleviate this effect, *τ* was adjusted to 50 ms, enhancing adaptation speed while preserving phasic-tonic behaviour. After optimization, there was a non-negligible MAE of 20 Hz. Moreover, our hair field afferents (Fig 3A) encoded considerably lower deflection angles than observed for the cockroach hair field (Fig 3C). Finally, our model is deterministic in that it does not include any noise.

By choosing to optimize the spike dynamics according to a proprioceptive hair plate of the American cockroach, we assumed that these dynamics would be consistent across different species and limbs. However, given the fact that even hairs within the same hair plate may differ with regard to details of their spike response [55] a general-purpose model needs to be tunable. Indeed, the AdEx model allows simple tuning of phasic peak and tonic steady-state spike rates by adjusting specific model parameters. To demonstrate this, the supplementary material shows how to adjust phasic and/or tonic responses as desired. For instance, varying *a* modifies *f*_ss_ while only minimally affecting the phasic peak (S3 Fig). Increasing *C* while decreasing *a* alters the strength of the phasic peak without changing *f*_ss_ (S4 Fig). Additionally, the AdEx model is capable of generating a purely tonic response by setting *b* to zero and modifying *C* (S5 Fig). Since most proprioceptors display a phasic and/or tonic response, this demonstrates the model’s versatility in simulating a wide range of proprioceptors and their dynamics.

We conclude that the AdEx model may serve a general template for spiking proprioceptor afferents with phasic-tonic response dynamics. To obtain an even more biologically accurate model, future studies could explore methods to account for variations in hair length, analyze possible nonlinear relationships between joint and hair angle, or optimize hair distribution. The latter could reduce redundancy and/or entail the implementation of the efficient coding hypothesis [56]. This hypothesis states that sensory systems are configured to efficiently encode the dynamic sensory stimuli that the organism encounters in its environment. Therefore, these systems are unlikely to adhere to a strict linear structure; instead, they are likely arranged to maximize coding efficiency for the most common sets of stimuli encountered by the organism. This concept can also be applied to the organization of the hair fields, suggesting that hairs can exhibit variation in receptive field size based on the probability distribution of joint positions.

### Layer two: Position interneurons (INs)

Position-sensitive second-order INs were modelled to extract tonic proprioceptive information from the hair field layer. The design drew inspiration from so-called ‘simple position-sensitive’ descending INs of the stick insect antennal system [19, 34]. Together with ‘dynamic position-sensitive’ descending INs that spike whenever the antenna is moving across a particular joint angle range, they are well-documented examples of position-encoding first-order INs in general. Position-dependent encoding of joint angles of foot position has also been found in thoracic INs of the stick insect [17, 57] but were mostly characterised with cyclic leg movement stimuli that combined position- and motion cues. Concerning the putative proprioceptors driving these IN responses, the trochanteral hair field has been shown to be the sole sensory input to the feedback control loop at the coxa-trochanter joint [58]. Ablations of this hair plate affects swing height during walking [39], the amplitude of searching movements [59].

Other than an earlier model by Ache and Dürr [34], the present study uses a spiking neuron model to implement position-encoding first-order INs. Consistent with our study, they found that employing two hair fields connected to two separate INs allowed to estimate joint angle reliably from a natural stimulus.

The non-zero MSE in the position layer (MSE = 0.030 55) arose due to two main factors: Overshoot (Fig 4D, *t* = 0.5, 1.5, 2.5) and stair-stepping (Figs 4C,D). Overshoot occurred due to the phasic peak in the sensory neuron activity during changes in joint angle direction. The position INs adopted this peak and, therefore, overshot relative to the experimental joint angle. The overshoot was exacerbated by increasing *ω*, as evidenced by the rise in MSE depicted in Fig 3B. To mitigate this overshoot, one could introduce a low-pass filter between each sensory neuron and its corresponding position IN. To do so, *f*_c_ should be set just above *f*_ss_, similar to the tuning of velocity INs. Stair-stepping arose as an artifact from the simulation time step and the calculation of the spike rates. This artifact can be reduced by shortening the time step or by smoothing the spike rate time course.

### Layer two: Velocity interneurons (INs)

As a complement to position INs, velocity INs were designed to extract the phasic component of afferent activity, and to spike only if the joint angle changed. Inspiration was drawn from ‘ON-type velocity sensitive’ INs in the stick insect antenna [19, 37], which receive proprioceptive input from pedicellar hair fields [60]. There is also evidence for velocity-sensitive INs in the antennal systems of crickets [20] and the fruitfly [4]. In comparison with an earlier model [34], our present model replaces filter blocks by a spiking neural network. As their physiological counterparts, our velocity INs demonstrated linearity between spike rate and angular velocity (Fig 5A) and precise encoding of angular velocity in response to realistic stimuli (Fig 5D, MSE = 0.3421). Velocity encoding was direction-selective (Figs 5B,C) with a high accuracy of 0.914.

Strong linearity was achieved only when *ω* equals 10.8 mV. At this synaptic strength, *f*_c_ marginally exceeded *f*_ss_, allowing spikes from small phasic fluctuations to transmit through the high-pass filter. However, for lower values of *ω, f*_c_ increased, resulting in complete blockage of spikes from small phasic fluctuations by the high-pass filter and zero postsynaptic spikes at low angular velocities (Fig 5A). Moreover, increasing the number of hairs per hair field *N*_h_ also enhanced encoding of low angular velocities. Doubling *N*_h_ (from 25 to 50) halved the receptive field size and doubled the rate of change of hair angles for a given change in joint angle. This amplified the phasic peak for low angular velocities. Thus, the optimal parameter set would be *ω* = 10.8 mV and *N*_h_ = ∞. However, due to a biological constraint on the number of hairs, *N*_h_ = 50 was considered the maximum number of hairs per hair field [26].

The apex of the phasic peak occurred when a hair reached maximum deflection (Fig 3D). Consequently, the second half of the phasic peak lagged maximum hair deflection. This resulted in a slight delay (≈ 0.025 s) in the velocity INs’ spike rate compared to the actual joint velocity. This delay was clearly evident in Fig 5B and D, and largely contributed to the accuracy of 0.914. Upon correction of this delay, by manually shifting the model response by 0.025 s, the MSE decreased from 0.2812 to 0.0910 (compared to 0.030 55 for the position INs). This implies that the velocity INs were less accurate than the position INs. Position INs were capable of detecting joint angle changes within the receptive field of a hair, whereas velocity INs only responded to changes from one receptive field to another. Consequently, the sensitivity of velocity INs was directly linked to the number of receptive field edges and therefore *N*_h_.

There is some discrepancy between the ON-type neurons found in the stick insect and our velocity IN implementation. Firstly, both hair fields had to be extended (Fig 2) to achieve sensitivity across the entire joint angle working range. Secondly, our velocity INs were directionally selective, unlike the ON-type neuron [37]. Ache and Dürr [34] had already suggested two ways by which a velocity IN could lose the directional selectivity of its afferent input (post-excitatory rebound; additional interneurons). A further option would be to fully connect all hairs from a extended bi-directional hair field (Fig 2B). Here, we renounced any of these options because direction-selectivity demonstrated distinct advantages in the subsequent network layers (see companion paper: van der Veen et al. [23]). Thirdly, the current network architecture solely relies on afferent input from hair fields. However, in insects, velocity information is also contributed to the proprioceptive system by the chordotonal organ [1, 9, 25]. It could be interesting to explore whether incorporating additional proprioceptors would mitigate the issues raised.

### Future work

We conclude that two fundamental aspects of distributed proprioceptive encoding of joint kinematics can be modelled by a two-layered SNN: phasic-tonic encoding and the extraction of joint angles and joint angle velocity at high accuracy. Future research may expand on these results, e.g., by tuning the AdEx neuron model parameters to different proprioceptor types (e.g., scolopidial neurons of chordotonal organs) [3, 4], by introducing multimodal integration among different proprioceptors [7], or by detailed models of distinct classes of first-order interneurons [19, 34, 57].

The companion paper [23] expands the present network by incorporating ‘movement primitive’ neurons that are designed to extract more complex intra-leg information about step cycle phases and/or transitions between them. Additionally, it introduces posture neurons that are capable of encoding higher-order information about whole-body posture from movement primitive neurons of all six legs. In combination with the present study, the proposed four-layered SNN links aspects of primary proprioception, inter-segmental proprioceptive pathways, and internal representation of whole-body posture.

## Supporting information

**S1 Fig The AdEx neuron dynamics**. The AdEx model response to a sustained current, I, governed by Eqs (6, 7, 8). This current is integrated into the membrane voltage, V. When *V* = *V*_T_ = −50 mV, the membrane voltage spikes and resets to *E*_L_ = −70 mV. During a spike event, 0.264 nA is added to *ω*, counteracting the input current. The model exhibits lower spike frequencies with increasing *ω* until reaching an equilibrium spike rate. The model parameters are given in Table 1 (sensory neuron).

**S2 Fig The LIF model dynamics**. The LIF model responds to a constant spike rate (333.3 Hz), governed by Eqs (9, 10, 11). A presynaptic spike, Pre, increases the membrane potential by 10 mV, and if no spikes are present, the membrane voltage, *V*, decays back to *E*_L_ = −70 mV. When *V* > *V*_T_ = −50 mV, a postsynaptic spike, Post, is recorded, and *V* resets to *E*_L_ = −70 mV. The postsynaptic spike rate remains constant in response to a constant presynaptic spike rate, since the LIF model has no adaptation. At the shown input spike rate, every third presynaptic spike triggers a postsynaptic spike. The model parameters are given in Table 1 (velocity neuron).

**S3 Fig The AdEx model response to varying** *a*. The AdEx model’s spike response to a ramp-and-hold hair deflection at an angular velocity of 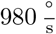 The parameters for the AdEx model were taken from Table 1. The parameter *a* was varied to demonstrate that the tonic steady-state frequency can be reduced while maintaining a strong phasic peak. A purely phasic response can be achieved by combining the AdEx model with a high-pass filter in series.

**S4 Fig The AdEx model response to varying** *C* **and** *a*. The AdEx model’s spike response to a ramp-and-hold hair deflection at an angular velocity of 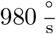 The parameters for the AdEx model were taken from Table 1. The parameters *C* and *a* were varied to demonstrate that the strength of the phasic peak can be adjusted without altering *f*_ss_.

**S5 Fig The AdEx model response to varying** *C*. The AdEx model’s spike response to a ramp-and-hold hair deflection at an angular velocity of 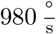 The parameters for the AdEx model were taken from Table 1, with *b* set to zero to eliminate adaptation and yield a purely tonic response. The capacitance *C* was adjusted to achieve different values of *f*_ss_.

## Acknowledgments

The authors would like to acknowledge the financial support of the CogniGron research center and the Ubbo Emmius Funds (Univ. of Groningen). Arne Gollin supported the project by sharing a curated motion capture dataset.

## Notes

### Competing Interest Statement

The authors have declared no competing interest.

